# The Big Five, Self-efficacy, and Self-control in Boxers

**DOI:** 10.1101/361295

**Authors:** Xin Chen, Bing Li, Jiaqiong Xie, Yun Li, Guodong Zhang

**Author notes:** Correspondence: Guodong Zhang.

## Abstract

Inviting 210 boxers of national athletes in China as participants, this study applied the NEO Five-Factor Inventory and self-control and self-efficacy scales for athletes to examine the relationship between personality traits and self-control, as well as any effect of self-efficacy as a mediator between the two variables. The data analysis indicated that, firstly, the boxers’ overall level of self-control is high, and the higher the competitive level, the higher the level of self-control. Secondly, there were significant correlations among the Big Five, self-control, and self-efficacy. Thirdly, the mediation model showed that self-efficacy has a significant mediating effect between the Big Five and self-control. These results suggest that formulating training and intervention programs based on the personality traits of boxers and focusing on training their self-efficacy (1) to help them enhance their self-control ability, thereby improving athletic performance and promoting physical and mental health, and (2) to support the inclusion of personality traits, self-efficacy, and self-control among psychological indicators to be assessed in boxers.

## Introduction

Self-control refers to the phenomenon of people overcoming their natural and automatic tendencies, desires, and behaviors and resist short-term temptations to achieve long-term goals [1]. It is essential for the well-being and healthy development of humans. Sigmund Freud believed that self-control is a major characteristic of a civilized society [2], while Hare et al. suggested that “the ability to exercise self-control is the key to human success and happiness” [3]. Good self-control not only prevents drug abuse, criminal offenses, and other undesirable social behaviors, but also promotes the healthy growth of individuals and the harmonious development of society [4,5]. Scholars believe self-control is the core element for achieving optimal competitive performance [6,7]. Boxers should utilize self-control, as it is a necessary quality [8,9]. Therefore, more emphasis should be put into research on the self-control of boxers. Scholars in sport psychology have called for research that “gives a voice” to marginalized groups, which would arguably include boxers [10,11].

Exploring the relationships between self-control, personality traits, and self-efficacy can serve as an entry point for the in-depth study of self-control. A report by Baniassadi et al. showed that self-control positively predicts responsibility and humanity, and negatively predicts neuroticism [12]. While Tianxin Mao et al. similarly find that openness, conscientiousness, extraversion, and agreeableness were all positively correlated with self-control, and neuroticism was negatively correlated with self-control [13]. In addition, Vera et al believes that self-control is affected by self-efficacy [14], and Studies et al. suggested that the higher the self-efficacy, the stronger the self-control ability [15]. In academic research, most of the existing research on self-control in sporting contexts focused on soccer players, divers, middle-distance runners, and college athletes of all types. At present, however, there is only limited research on the psychology of boxing in China, especially on the relationship between personality traits and self-control in boxers. Thus, this study invited Chinese boxers of national athletes to take part in an examination of the relationship between their personality traits and self-control, and to look for any mediating effect of self-efficacy between them. In doing so, the study clarifies the value of personality traits and the significance of the ability of self-control to help boxers improve their athletic performance and psychological indicators and to maintain physical and mental health.

### Personality traits and self-control

Most studies have shown that there is a significant correlation between the Big Five and self-control. Neuroticism has a negative correlation with self-control; agreeableness, extroversion, openness, and responsibility are positively related to self-control; self-control is a prerequisite for individuals to adapt to their social environment [16]. Besides, it had been found that there were close links between the Big Five and self-control in many studies, such as, a significant negative correlation was observed between neuroticism and self-control, and both agreeableness and conscientiousness were found significantly positively associated with self-control [17,18,19]. In addition, different researchers found differences in the relationship between extroversion and openness and self-control, some studies have found that extraversion and self-control had significant negative correlation [17,20], while some other studies found no significant correlation between them [18,19,21]. As to the relationship between openness and self-control, some studies found that there was a significant positive correlation [19], while some studies found no significant correlation [18,20,21]. Due to its consistency and stability across languages and cultures [22], the Big Five are often used to predict self-control [23]. Therefore, based on previous studies, this study proposes hypothesis 1: Neuroticism is negatively correlated with self-control, while agreeableness, conscientiousness, and extraversion are positively correlated with self-control.

### Personality traits and self-efficacy

Previous studies show that there is a significant correlation between the Big Five and self-efficacy: neuroticism is negatively correlated with self-efficacy, and extraversion, openness, agreeableness, and responsibility positively correlated with self-efficacy. Personality traits are an important factor influencing the self-efficacy of individuals [24]. Studies have linked the Big Five traits and self-efficacy [25, 26, 27]. Some scholars found that individuals with higher scores of conscientiousness had higher self-efficacy beliefs [28,29]. Openness shifts perceptions of demands into challenges to be tackled, broadening task engagement and self-efficacy [30]. Research has found agreeableness can lead to increased self-efficacy [31]. Ou et al. found that self-efficacy is positively correlated with conscientiousness, agreeableness, openness and extraversion, but negatively correlated with neuroticism [32]. Certain researchers have found that individual self-efficacy is positively correlated with extroversion and negatively correlated with neuroticism [33, 25]. The finding of Gordana et al. indicated that conscientiousness predicts the self-efficacy of teachers [34], while Marcionetti believes that conscientiousness, neuroticism, and extraversion are significantly correlated with self-efficacy [35]. Furthermore, Brown et al. proposes that higher conscientiousness and extraversion, and lower neuroticism, help enhance self-efficacy [36]. In sports, Wang et al. found that neuroticism has a significant negative predictive effect on the general self-efficacy of basketball players, while extraversion and conscientiousness have significant positive predictive effects [37]. Based on the existing literature, this study proposed hypothesis 2: neuroticism is negatively correlated with self-efficacy, while agreeableness, conscientiousness, and extraversion are positively correlated with self-efficacy in boxers.

### Self-control and self-efficacy

Based on previous studies, there is a positive correlation between self-control and self-efficacy. Bandura’s self-efficacy and self-regulation theories suggest that self-control is affected by self-efficacy, and there was a significant positive correlation between the two [38]. Many studies have found a positive correlation between self-efficacy and self-control. Graham et al. studied self-efficacy as a psychological factor to explain how self-control is performed [39]. Baumeister’s analysis showed that self-control requires an individual’s own control resources, and self-efficacy complements this resource by acting as a positive emotion [40]. Yu et al.’s study concluded that there is an interaction effect between self-efficacy beliefs and self-control behavior [41]. A large number of scholars have shown that there is a positive correlation between self-efficacy and self-control in different population groups [42,43]. Feldman et al. found that self-efficacy positively predicts self-control [44] in child. Jones et al. found that self-efficacy and sense of control are important indicators of an athlete’s state [45]. Li and Zhang also believe that self-efficacy positive had a positive effect on self-control in athletes [46]. However, does this mean that self-efficacy has a positive effect on self-control in boxers? Currently, there is a lack of research studies that address this issue. Therefore, this study proposed hypothesis 3: The self-efficacy and self-control of boxers are positively correlated.

### The mediating effect of self-efficacy

By integrating self-regulation theory, self-efficacy theory, and cognitive emotion theory, we can observe that personality traits both directly affect self-control and indirectly influence self-control through self-efficacy. Some scholars emphasize the important role of self-efficacy in sports: it has a great impact on the performance and self-assessment of athletes [47] and is an underlying cause of athletic performance [48]. Several studies in the context of China have confirmed that self-efficacy has significant mediating effects in different research fields and between different personality variables. For example, the results of Huang et al. [42] and Xu et al. [49] indicated that self-efficacy has a partial mediating effect on the relationship between self-enhancement goal orientation and self-control, and a mediating effect between personality and psychological health. Fang et al. [43] found that self-efficacy plays a significant mediating role between the self-supporting personality of college students and self-control. Wang et al. found that self-efficacy has a mediating effect between personality traits and direct stress coping styles in female basketball players [37]. Thus, as a typical characteristic of excellent boxers, does self-efficacy have a mediating effect between personality traits and self-control? To find out, this study proposed hypothesis 4: the self-efficacy of boxers mediates the effects of neuroticism, agreeableness, conscientiousness, and extraversion on self-control.

## Subjects and Methods

### Participants

This study adopted cluster sampling and selected boxers from Chinese national boxing team and boxing teams from Sichuan, Chongqing, Guizhou, and Yunnan provinces as participants to complete a survey questionnaire. A total of 230 questionnaires were distributed, and 210 valid ones were returned; the effective recovery rate was 91%. Among the participants, 76 were men (36.2%) and 134 were women (63.8%). There were 79 Level 3 athletes (37.6%), 24 Level 2 athletes (11.4%), 49 Level 1 athletes (23.3%), 45 Master Level athletes (21.4%), and 13 athletes at the International Master Level (6.2%). Level 3 is the lowest and the International Master Level is the highest [50]. Their average age was 18.89 years (SD = 3.83) and their average prior training period was 4.93 years (SD = 3.22).

### Procedure

The participants in this study were all boxers of national athletes who were strictly trained before the survey was conducted. After receiving informed consent from the management, coaches, and athletes of the national team and other sports teams, the questionnaire was distributed to teams at the provincial level or above. The instructions were explained in detail and example questions were provided to the participants, who were asked to read the questionnaire carefully and answer it according to their actual circumstances. Between June 12 and July 8, 2017, 230 questionnaires were distributed at Chinese boxing team, Qujing in Yunnan province, Qingzhen in Guizhou province, Shapingba in Chongqing city, and Liangshan in Sichuan province. The questionnaires were collected on site and 210 were considered valid, achieving a 91% effective rate.

### Measures

This study employed the survey research method. Specifically, the following three questionnaires were included in the study.

### NEO Five-Factor Inventory (NEO-FFI)

This study used the NEO-FFI, created by psychologists McCrae and Costa [51]. The confirmatory factor analysis (CFA) results were good: *χ*2/*df* = 1.330, RMSEA = 0.04, TLI = 0.989, GFI = 0.979, NFI = 0.978, CFI = 0.994, IFI = 0.994. The factor loadings of the items were between 0.50 and 0.73, indicating that the questionnaire has good structural validity. The NEO-FFI contains 60 items; item scores are added together to get a total score. The Cronbach’s alpha coefficients of the inventory’s five dimensions were 0.77 (Neuroticism), 0.76 (Agreeableness), 0.80 (Conscientiousness), 0.72 (Extraversion), and 0.57 (Openness). Openness is not consistent with the personal characteristics of boxers under Chinese cultural background, and in this study, the openness dimension reliability does not meet the requirements of the research standards, so the openness is deleted.

### Self-control questionnaire for athletes

This study used a self-control questionnaire for athletes created by the Chinese scholars Li Xiaoyu and Zhang Liwei [52]. The self-control ability of boxers was assessed, and the CFA results were good: *χ*^2^/*df* = 1.148, RMSEA = 0.027, TLI = 0.971, GFI = 0.916, NFI = 0.865, CFI = 0.979, IFI = 0.980. The factor loadings of the items were between 0.49 and 0.75, indicating the questionnaire has good structural validity. The questionnaire contains 24 items in total, and the item scores are added together to get a total score. The higher the total score, the better the self-control of the individual. The Cronbach’s alpha coefficient of the questionnaire was 0.87.

### Self-efficacy scale for athletes

This study used a self-efficacy scale for athletes created by the Chinese scholars Wei Ping and Chen Hongbo [53]. The questionnaire’s applicability to boxers was tested, and the CFA results were good: *χ*^2^/*df* = 1.136, RMSEA = 0.025, TLI = 0.990, GFI = 0.956, NFI = 0.950, CFI = 0.994, IFI = 0.994. The factor loadings of the items were between 0.43 and 0.75, indicating good structural validity. The questionnaire contains 15 items, and the item scores are added together to get a total score. The higher the total score, the higher the level of self-efficacy of the individual. The Cronbach’s alpha coefficient of the questionnaire was 0.92.

### Analyses

This study used the bias-corrected percentile bootstrap method recommended by Fang and Zhang (2012) [54] in the significance testing of all regression coefficients. To implement this method, we used the Model 4 (which hypothesizes that a mediating effect exists, consistent with the theoretical hypothesis model of this study) PROCESS macro for SPSS created by Hayes (2012) [55] and tested the hypothesized models by controlling for gender, age, years of exercise, and competitive level and estimating the 95% confidence interval of the mediating effect through 5,000 samples.

## Results

### Testing for common method bias

In this study, some items in the questionnaires were expressed in reverse. Moreover, all questionnaires were anonymous, and the experimental procedures addressed the risk of common method bias in collecting data using questionnaires thus, through a method proposed by previous researchers [56]. That is, using AMOS 21.0, the common factor of all variables was set to 1, and all items variable was used as explicit variables to conduct CFA. The CFA results showed that the model fit index (*χ*^2^/*df* = 2.008, RMSEA = 0.069, NFI = 0.337, CFI = 0.498, TLI = 0.486, GFI = 547, IFI = 0.504) was low, indicating no serious common method bias.

### Analysis of self-control and self-efficacy among Chinese boxers

On a five-point Likert scale, the mean score of self-control was M = 3.68, indicating that the overall level of self-control among boxers in China is high. This study also examined gender differences and competitive level differences in self-control; the results indicated no significant gender differences (*F* = 1.14, *P* = 0.28), but a significant major effect of competitive level (*F* = 7.81, *P* = 0.00). The interaction between gender and competitive level was not significant (*F* = 1.82, *P* = 0.13). The mean self-control scores of boxers from the five different competitive levels were significantly different: the higher the competitive level, the higher the level of self-control (International Master Level M = 3.92; Master Level M = 3.79; Level 1 athletes, M = 3.77, Level 2 athletes, M = 3.83; Level 3 athletes, M = 3.47). The mean score of self-control in International Master Level is higher than that of the Level 3 (*P* < 0.01).

The mean score of self-efficacy was M = 3.50, indicating that the overall score of self-efficacy among boxers in China is high. In terms of the level of self-efficacy, there was no significant difference between male and female boxers (*P* > 0.05). The mean score of self-efficacy among boxers from five different competitive levels were significantly different: the higher the competitive level, the higher the self-efficacy score (International Master Level M = 3.81; Master Level M = 3.66; Level 1 athletes, M = 3.53, Level 2 athletes, M = 3.60; Level 3 athletes, M = 3.30). There was a significant difference on self-efficacy between International Master Level and Level 3 (*P* < 0.01), too.

### Correlation analyses of personality traits, self-efficacy, and self-control

The correlation analyses (Table 1) show that neuroticism is significantly and negatively correlated with self-efficacy and self-control, while extraversion, agreeableness, and conscientiousness are significantly and positively correlated with self-efficacy and self-control. And more, self-efficacy and self-control are positively correlated. These significant correlations between variables provided a good basis for subsequent testing for mediating effects, while confirming H1, H2, and H3.

**Table 1.** Means, standard deviations, and correlation coefficients of personality traits, self-control and self-efficacy.

|  | M | SD | 1 | 2 | 3 | 4 | 5 | 6 |
| --- | --- | --- | --- | --- | --- | --- | --- | --- |
| **Neuroticism** | 2.94 | 0.53 | 1 |  |  |  |  |  |
| **Extraversion** | 3.63 | 0.49 | -0.399*** | 1 |  |  |  |  |
| **Agreeableness** | 3.78 | 0.46 | -0.492*** | 0.371*** | 1 |  |  |  |
| **Conscientiousness** | 3.50 | 0.48 | -0.519*** | 0.470*** | 0.537*** | 1 |  |  |
| **Self-efficacy** | 3.50 | 0.64 | -0.416*** | 0.481*** | 0.448*** | 0.698*** | 1 |  |
| **Self-control** | 3.68 | 0.49 | -0.566*** | 0.476*** | 0.677*** | 0.741*** | 0.704*** | 1 |

Note: \* indicates $P < 0.05$ , \*\* indicates $P < 0.01$ , and \*\*\* indicates $P < 0.001$

### Testing for mediation by self-efficacy on effects of neuroticism, agreeableness, extraversion, and conscientiousness on self-control

This study used the Bootstrap method proposed by Fang et al. [54] and the Model 4 PROCESS macro for SPSS created by Hayes [55] to conduct mediating effect testing.

Regression analysis results (Tables 2-5) show that neuroticism significantly and negatively predicted self-efficacy (*β*= −0.23, *P* < 0.01) while self-efficacy significantly and negatively predicted self-control (*β*= 0.88, *P* < 0.001), as did neuroticism (*β* = −0.32, *P* < 0.001). Extraversion had a significant positive predictive effect on self-efficacy (*β*= 0.17, *P* < 0.001), while self-efficacy significantly and positively predicted self-control (*β*= 0.78, *P* < 0.001). Extraversion and self-efficacy both positively predicted self-control, extraversion significantly (*β*= 0.27, *P* < 0.001). Agreeableness significantly and positively predicted self-efficacy (*β*= 0.26, *P* < 0.001), which significantly and positively predicted self-control (*β*= 0.77, *P* < 0.001), as did agreeableness (*β*= 0.44, *P* < 0.001). Conscientiousness significantly and positively predicted self-efficacy (*β*= 0.43, *P* < 0.001), and self-efficacy significantly and positively predicted self-control (*β*= 0.58, *P* < 0.001), as did conscientiousness (*β*= 0.47, *P* < 0.001).

**Table 2.**
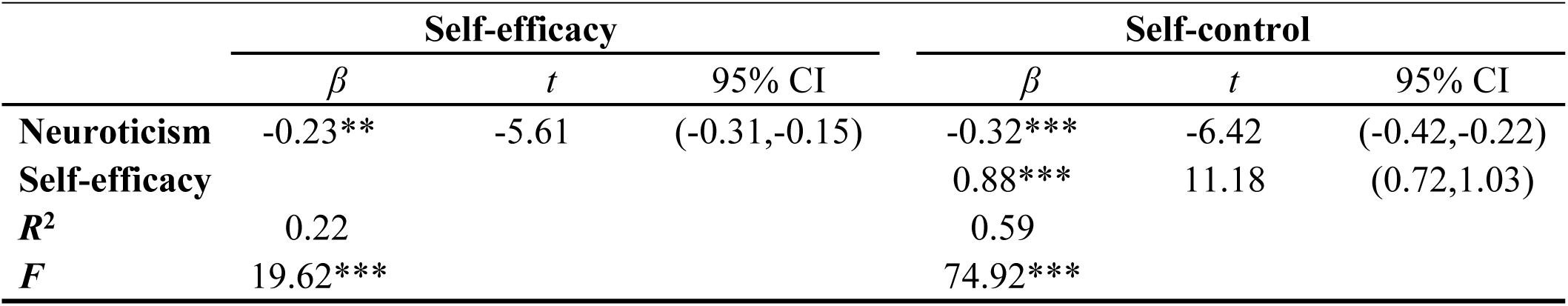
Regression analysis of the mediating effect of self-efficacy between neuroticism and self-control.

**Table 3.** Regression analysis of the mediating effect of self-efficacy between extraversion and self-control.

|  | Self-efficacy |  |  | Self-control |  |  |
| --- | --- | --- | --- | --- | --- | --- |
| | $\beta$ | $t$ | 95% CI | $\beta$ | $t$ | 95% CI |
| Extraversion | 0.17*** | 6.96 | (0.20,0.35) | 0.27*** | 3.14 | (0.06,0.28) |
| Self-efficacy |  |  |  | 0.78*** | 10.83 | (0.77,1.03) |
| $R^2$ | 0.27 | | | 0.53 | | |
| $F$ | 25.94*** | | | 58.83*** | | |

**Table 4.** Regression analysis of the mediating effect of self-efficacy between agreeableness and self-control.

|  | Self-efficacy |  |  | Self-control |  |  |
| --- | --- | --- | --- | --- | --- | --- |
| | $\beta$ | $t$ | 95% CI | $\beta$ | $t$ | 95% CI |
| Agreeableness | 0.26*** | 6.68 | (0.19,0.35) | 0.44*** | 9.41 | (0.35,0.54) |
| Self-efficacy |  |  |  | 0.77*** | 10.39 | (0.62,0.91) |
| $R^2$ | 0.26 | | | 0.66 | | |
| $F$ | 24.53*** | | | 99.19*** | | |

**Table 5.**
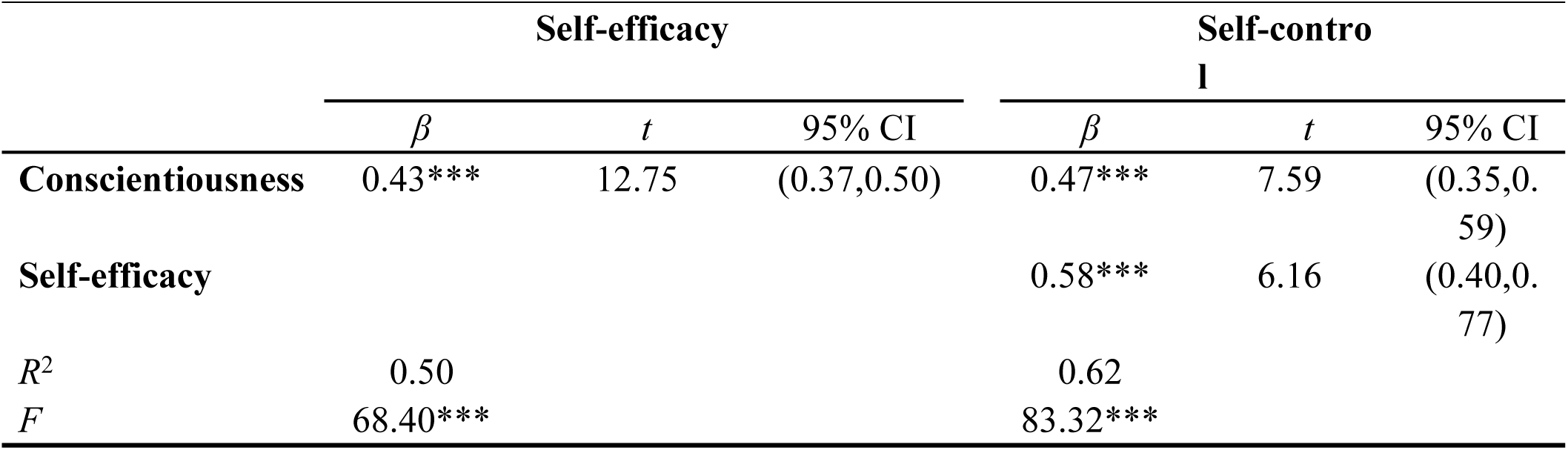
Regression analysis of the mediating effect of self-efficacy between conscientiousness and self-control.

Figs 1-4 present the standardized effect sizes and significance results corresponding to the effect paths of neuroticism, extraversion, agreeableness, and conscientiousness on self-control. The bootstrap 95% confidence interval of the indirect effect of self-efficacy does not contain a value of 0, indicating that self-efficacy has a significant mediating effect on the effects of neuroticism, extraversion, agreeableness, and conscientiousness on self-control. There are four mediation effect models outlined below, one for each personality trait variable. Taking the results all together, neuroticism explained 22% of the change in self-efficacy, extraversion 27%, agreeableness 26%, and conscientiousness 50%. As such, H4 was confirmed.

**Fig. 1.**
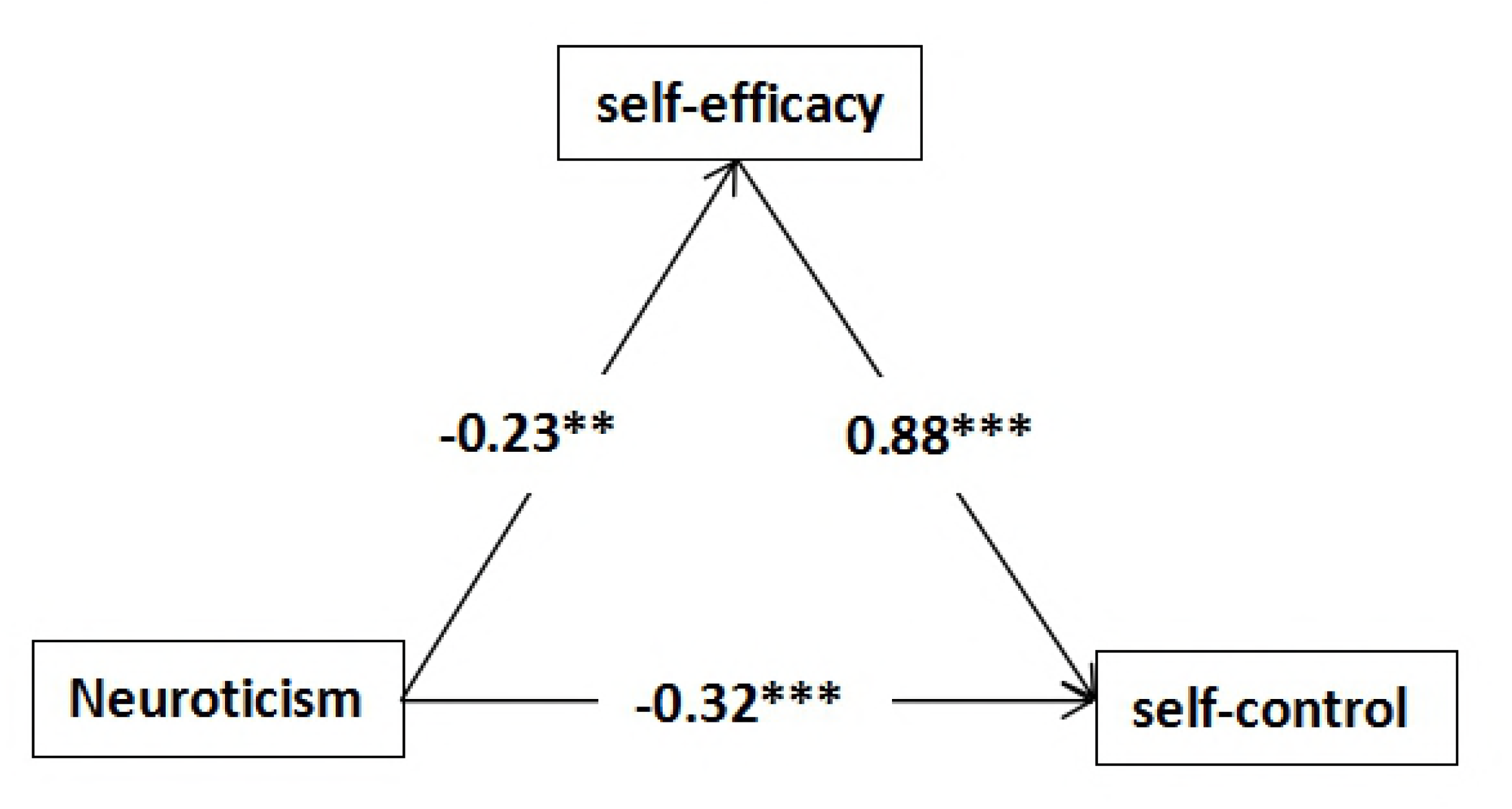
Neuroticism → self-efficacy → self-control.

**Fig 2.**
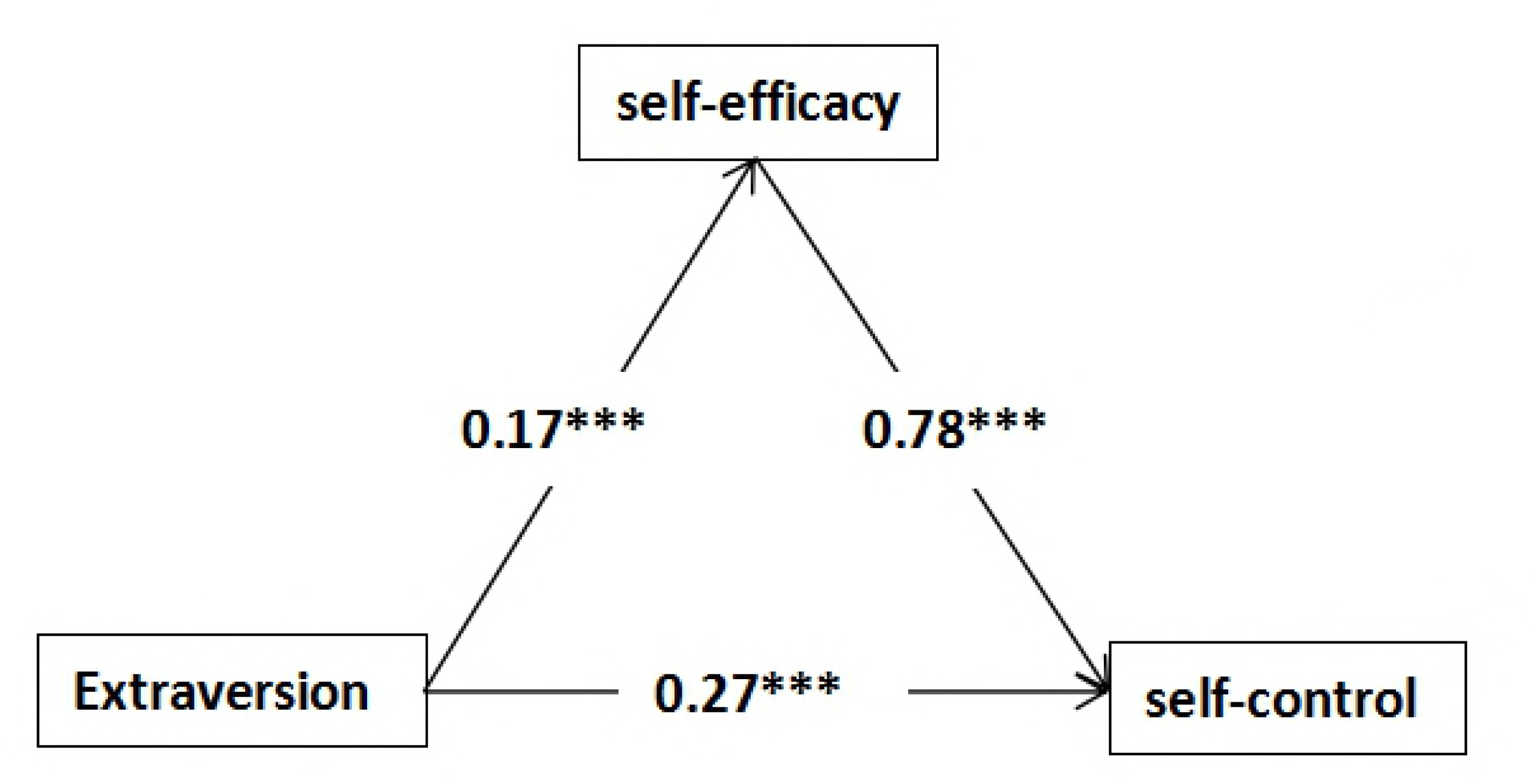
Extraversion → self-efficacy → self-control.

**Fig 3.**
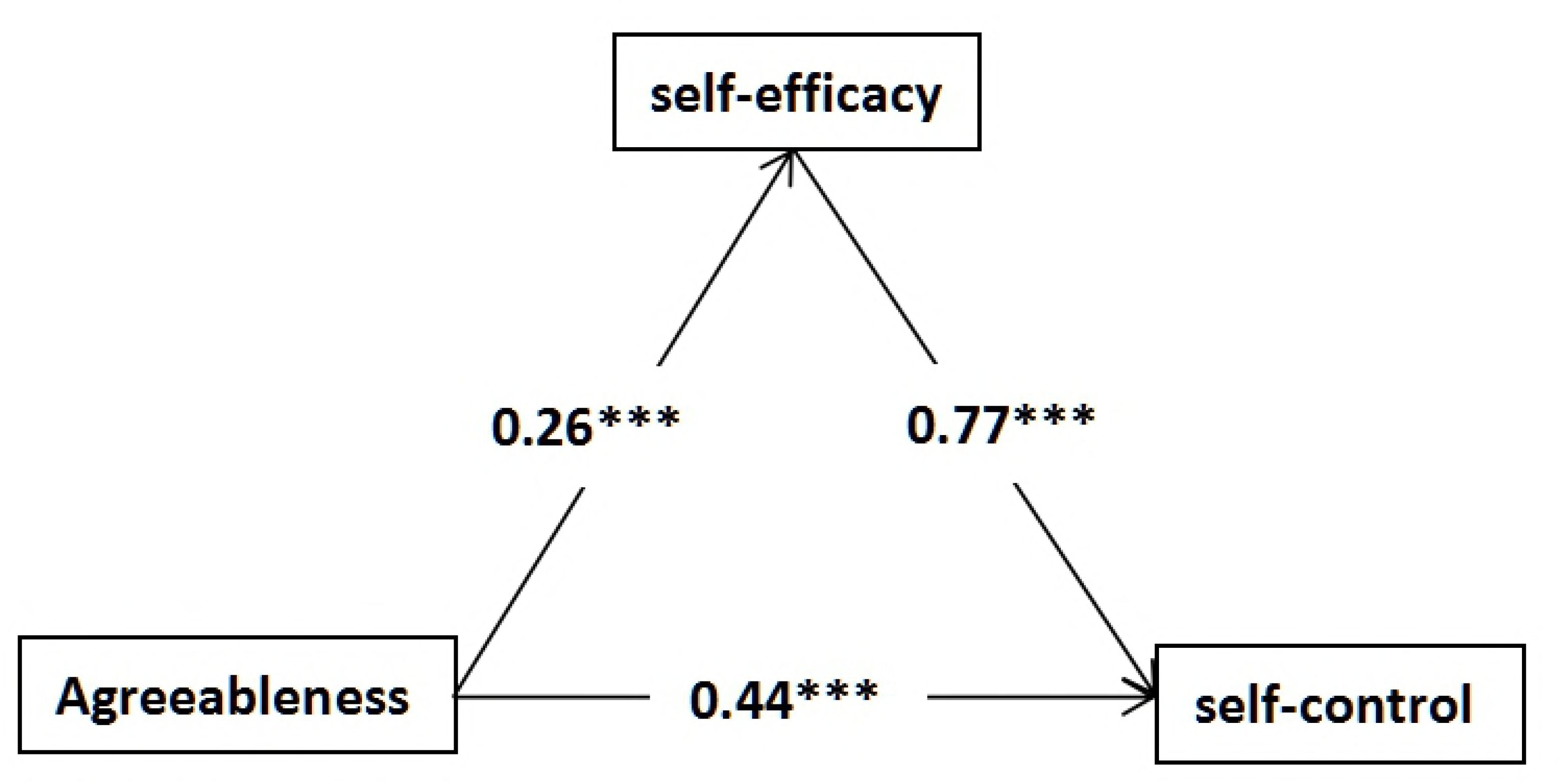
Agreeableness → self-efficacy → self-control.

**Fig 4.**
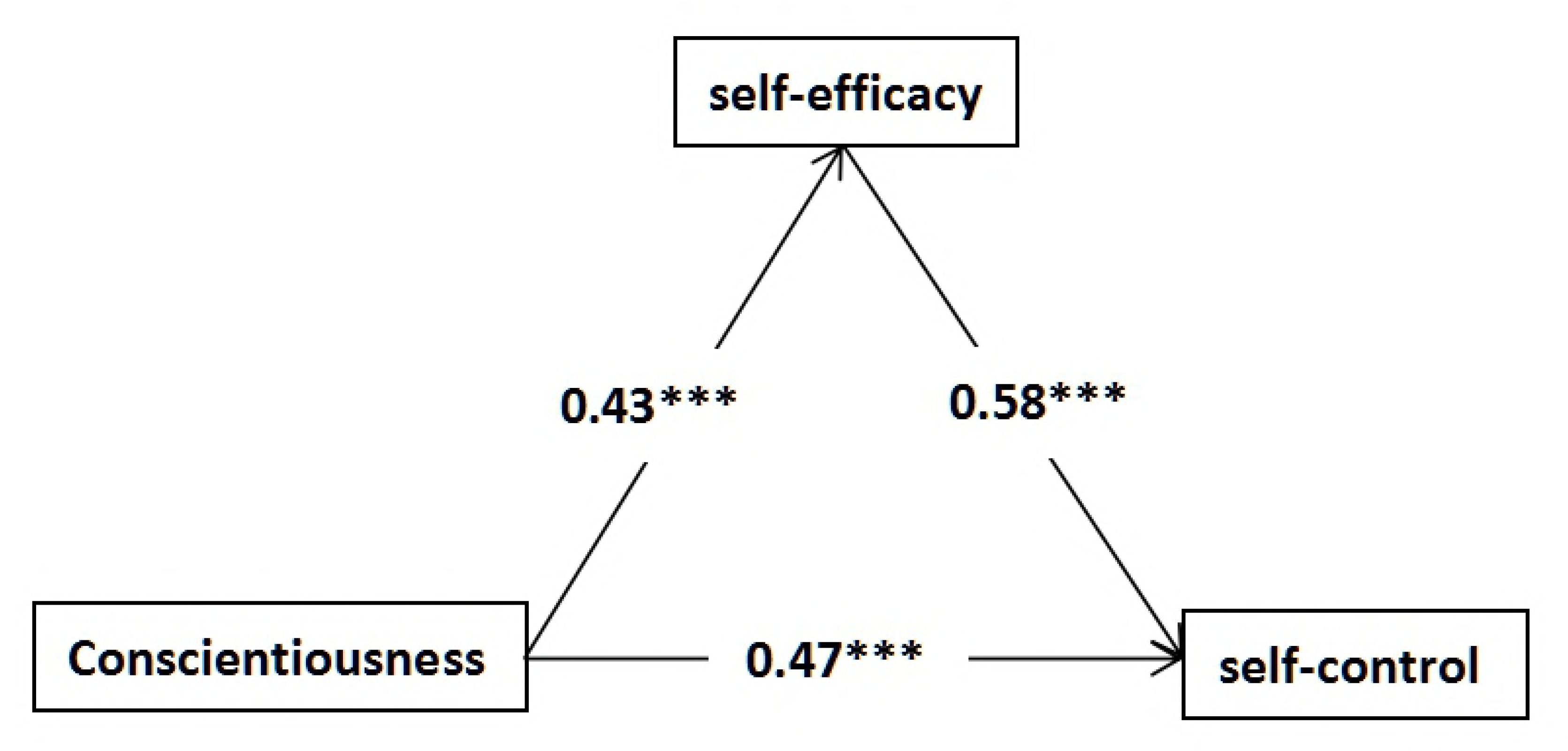
Conscientiousness → self-efficacy → self-control.

## Discussion

### Direct influence of personality traits on self-control

Chris believes self-control is an important factor in various sporting and training activities as well as in the achievement of excellent athletic performance [7]. It is also a key factor in reducing an individual’s fear of personal failure [57]. Zhou found that the ability of boxers to self-control, especially anxiety, has a direct influence on their use of skills and tactics [58]. Previous studies have indicated that agreeableness, conscientiousness, and extraversion are significantly and positively correlated with self-control [18,20], while neuroticism is significantly and negatively correlated with self-control [19,21]. Therefore, investigating the effect mechanism between personality traits and self-control may help practitioners study self-control in athletes from a personality trait perspective.

Based on previous research, this study hypothesized that neuroticism is negatively correlated with self-control, while agreeableness, conscientiousness, and extraversion are positively correlated with self-control. These hypotheses were supported by the results of data analysis and are consistent with the results of Bazzy [18], Deng et al. [59], and Pilarska [22]. Boxers can not only avoid injuries but also reduce the number of fouls by improving their self-control level and their own skills and tactics. [60]. Song et al. suggested that boxers should continuously enhance their psychological stability by focusing on the actual combat element of boxing [61]. Therefore, the relationship between the Big Five and the self-control of boxers has a certain significance in guiding the psychological selection and training of future boxers.

### The mediating role of self-efficacy between personality traits and self-control

Studies of the relationship between personality traits and self-control have shown that four of the Big Five have indirect predictive effects on self-control through self-efficacy. Theories of self-efficacy hold that it is the result of measuring and evaluating one’s own abilities [38]—the subjective judgment and self-feeling that individuals have about their ability to perform a certain behavior or task—and that it plays an important role in the decision-making process before a behavior is performed [62]. In sporting contexts, scholars have found that the stronger athletes’ self-efficacy, the better their performance in sports as diverse as athletics, tennis, scuba diving, and gymnastics [47], because when athletes perform a task for the first time, their self-efficacy affects their performance [63]. Other studies have demonstrated that, even despite excellent skills and motivation to win, people with outstanding capabilities are less likely to achieve success if they have poor self-efficacy [64]. The present study hypothesized that neuroticism is negatively correlated with self-control, while agreeableness, conscientiousness, and extraversion are positively correlated with self-control; and these hypotheses were supported by the results of data analysis and consistent with the results of Zhang et al. [65], Ou et al. [32], and Marcionetti [36]. Moreover, this study found that self-efficacy has a significant positive predictive effect on self-control, consistent with the results of Huang et al. [42], Li et al. [46], and Kaida et al. [66]

This study also hypothesized that self-efficacy mediates the effects of neuroticism, agreeableness, conscientiousness, and extraversion on self-control. Data analyses indeed revealed four mediating paths, showing partial mediating effects of self-efficacy on self-control for each personality trait variable. Many other researchers have also found self-efficacy have a mediating effect between personality traits and self-control, such as Wang et al. [37], Fang et al. [43], and Alexander et al. [67]. Therefore, cultivating and upgrading the self-efficacy of boxers should be emphasized in their training.

## Conclusions and Suggestions

### Conclusions

1. Overall, level of boxers’ self-control was high. There were significant differences in self-control between boxers of different competitive levels, and the higher the competitive level, the higher the self-control ability score. The same goes for self-efficacy.
2. There are significant correlations of neuroticism and agreeableness, conscientiousness, and extraversion with self-control, indicating that these four dimensions have direct predictive effects on self-control. Self-efficacy is positively correlated with self-control.
3. Boxers’ self-efficacy mediates between personality traits and self-control, indicating that personality traits predict self-control not only directly but also
indirectly through self-efficacy. Overall, the mediation effect models constructed in this study had high explanatory power for self-control.

## Suggestions

1. Based on the relationship between the Big Five and self-control in boxers, training and intervention programs can be adjusted according to the personality traits of specific boxers to improve their self-control ability.
2. The role of psychological indicators should be emphasized, and boxers’ psychological indicators improved by considering personality traits, self-efficacy, and self-control as key indicators.
3. The mediating effect of self-efficacy in boxers should be cultivated through daily training and competitions to enhance their self-control ability, ultimately helping them improve their performance.

## Acknowledgments

XC, BL, JQX, YL, and GDZ have satisfied all the criteria for authorship: substantially contributing to the study’s conception and interpretation; drafting and revising the work; approving the version to be published; agreeing to be accountable for the work.

